# Excitatory neurons of the anterior cingulate cortex encode chosen actions and their outcomes rather than cognitive state

**DOI:** 10.1101/2024.04.12.589244

**Authors:** Martin M. Jendryka, Uwe Lewin, Sampath K.T. Kapanaiah, Hartmut Dermutz, Birgit Liss, Anton Pekcec, Thomas Akam, Benjamin F. Grewe, Dennis Kätzel

## Abstract

The anterior cingulate cortex (ACC) causally influences cognitive control of goal-directed behaviour. However, it is unclear whether ACC directly encodes cognitive variables like attention or impulsivity, or implements goal-directed action selection mechanisms that are modulated by them. We recorded ACC activity with miniature endoscopic microscopes in mice performing the 5-choice-serial-reaction time task, and applied decoding and encoding analyses. ACC pyramidal cells represented specific actions before and during the behavioural response, whereas the response type (e.g. correct/incorrect/premature) – indicating the state of attentional and impulse control – could only be decoded during and after the response with high reliability. Devaluation and extinction experiments further revealed that action encoding depended on reward expectation. Our findings support a role for ACC in goal-directed action selection and monitoring, that is modulated by cognitive state, rather than in tracking levels of attention or impulsivity directly in individual trials.

## Introduction

Maintenance of high levels of sustained attention and inhibition of impulsive responding are key to successful goal-directed behaviour, and impaired in a variety of psychiatric disorders [1,2]. Both aspects can be measured by the 5-choice-serial-reaction-time task (5-CSRTT) [3,4] in both humans and rodents. In this task, subjects can make four different types of response, indicative of different cognitive states: (*i*) they can correctly respond to a stimulus presented briefly and after a considerable waiting time to earn a reward (correct response), requiring high attentional and impulse control. (*ii*) they can, instead, follow the impulsive urge to respond before cue-presentation (premature response, indicating reduced impulse control), or (*iii*) respond into a non-cued hole (incorrect response, indicating reduced attentional control). (*iv*) Alternatively, they may not respond at all (omission, indicating reduced task engagement or inattention). Therefore, the measurement of neurophysiological correlates of these four response options, promises to identify circuits that regulate attention, impulse control, and possibly other aspects of deterministic goal-directed behaviour.

Several rodent studies have implicated the anterior cingulate cortex (ACC) in this regulation. Manipulations of rodent ACC have been shown to produce shifts in the relative occurrence of these behavioural outcomes in the 5-CSRTT which support a causal role of this brain structure. For example, the activation of Gi-protein signalling in excitatory pyramidal cells, either in all layers or in layer 5 exclusively, may reduce premature and, partly, increase correct responding [5]. In contrast, the chemogenetic inhibition of a subgroup of ACC neurons projecting to the visual cortex may induce a shift from correct responding to response omission [6], whereas their pre-cue stimulation at 30 Hz after such errors may have the opposite effect [7]. The chemogenetic activation of ACC parvalbumin interneurons, in turn, reduces both premature and incorrect responses, but not response omissions [8].

Studies with physiological measurement of neural activity in rodent ACC during the 5-CSRTT and related tasks, have partly supported the possibility of such a causal role. One study revealed that excitatory and inhibitory neurons in rat ACC may change their firing rate differently both before and after correct vs. incorrect choices in the 5-CSRTT [9]. Specifically, ramping neural activity in the ACC and the adjacent prelimbic cortex (PrL, upper part of the rodent medial prefrontal cortex, mPFC) before cue-presentation has been interpreted as preparatory signal under conditions that require high sustained attention, as this activity increase was smaller before incorrect (low attention) responses and lowest during omissions [9,10].

Several other studies using physiological measurements have, however, failed to find such an indication of a causal role of ACC for modulating the occurrence of response options on a trial-by-trial basis in the form of distinct pre-choice activity. They rather suggest that ACC may monitor ongoing behaviour, and potentially provide feedback or error signals. For example, neurons in rat ACC were shown to encode behaviour-related information mostly during and after a choice, in a deterministic lever-based working memory task, thereby monitoring action and outcome [11]. A specific subpopulation of ACC neurons that project to visual cortex was selectively excited *after* incorrect choices or omissions (i.e. they conduct error-monitoring), but their activity did not differ between those erroneous and correct choices while they were made [7]. Imaging during a head-fixed Go/No-Go paradigm even found no evidence for a selective recruitment of these neurons and projections for enhanced stimulus discrimination, but rather that they simply represent rewarded action and stimuli [12]. Using a Go/No-go paradigm with visual cues in mice, another group confirmed that ACC neurons are generally more likely activated by cues that imply reward than those that do not, but also suggested that these cells fire selectively either to signals that imply action or action restraint [13]. Another study in the 5-CSRTT also failed to detect much increase of firing rates of pyramidal neurons in the dorsal PrL/ACC region before cue-onset, but found the modulation of their firing times by gamma-oscillations in this period [14]. The role of the ACC may also depend on the task structure, as it was shown that, in a probabilistic task, rat ACC neurons represent expected outcome first, before switching to actual outcome in case of a mismatch between the two [11], which could constitute a feedback signal for updating prior believes. This is in line with selective activity of some ACC neurons after incorrect choices in the 5-CSRTT, constituting an error signal [7,9].

In summary, no clear picture of the mechanistic role of the rodent ACC in attention and impulse control has emerged yet; whereas some studies found representations several seconds before the choice event, which could indicate a causal role for choices or, generally, the present cognitive state, others have found representations rather around the time of choice itself and most profoundly during rewarded or after incorrect responses, which is more in line with action- and outcome-monitoring, at least in deterministic tasks. Likewise, in monkeys, ACC activity has been linked to error-monitoring, value representation and belief-updating [15,16] rather than to attention *per se* [12]. Therefore, we here use simultaneous monitoring of dozens of excitatory neurons with miniature endoscopic microscopes (miniscopes) in the 5-CSRTT, in mice, in combination with time-resolved encoding and decoding analysis to reveal which aspects of attentional, impulse and motor control are represented in the ACC at which point in time.

## Results

### Miniscope-based recording of neocortical activity in the 5-CSRTT

To monitor activity of individual pyramidal neurons, we transduced ACC with an AAV5-vector expressing the fluorescent calcium sensor GCaMP6m under the CamKIIα-promoter [17], in male C57BL/6J wildtype mice (*N* = 12), and implanted a gradient refractory index lens in a separate surgery at the same location (Figure 1B). For comparison, we also generated a smaller, second subgroup (*N* = 6), where activity was monitored in the ventral mPFC (Figure 1B), a region that was previously shown to represent rewarded choices [18]. Mice had been pre-trained in the 5-CSRTT, and their training was continued after recovery from the second surgery, until they reached a stable baseline.

**Figure 1.**
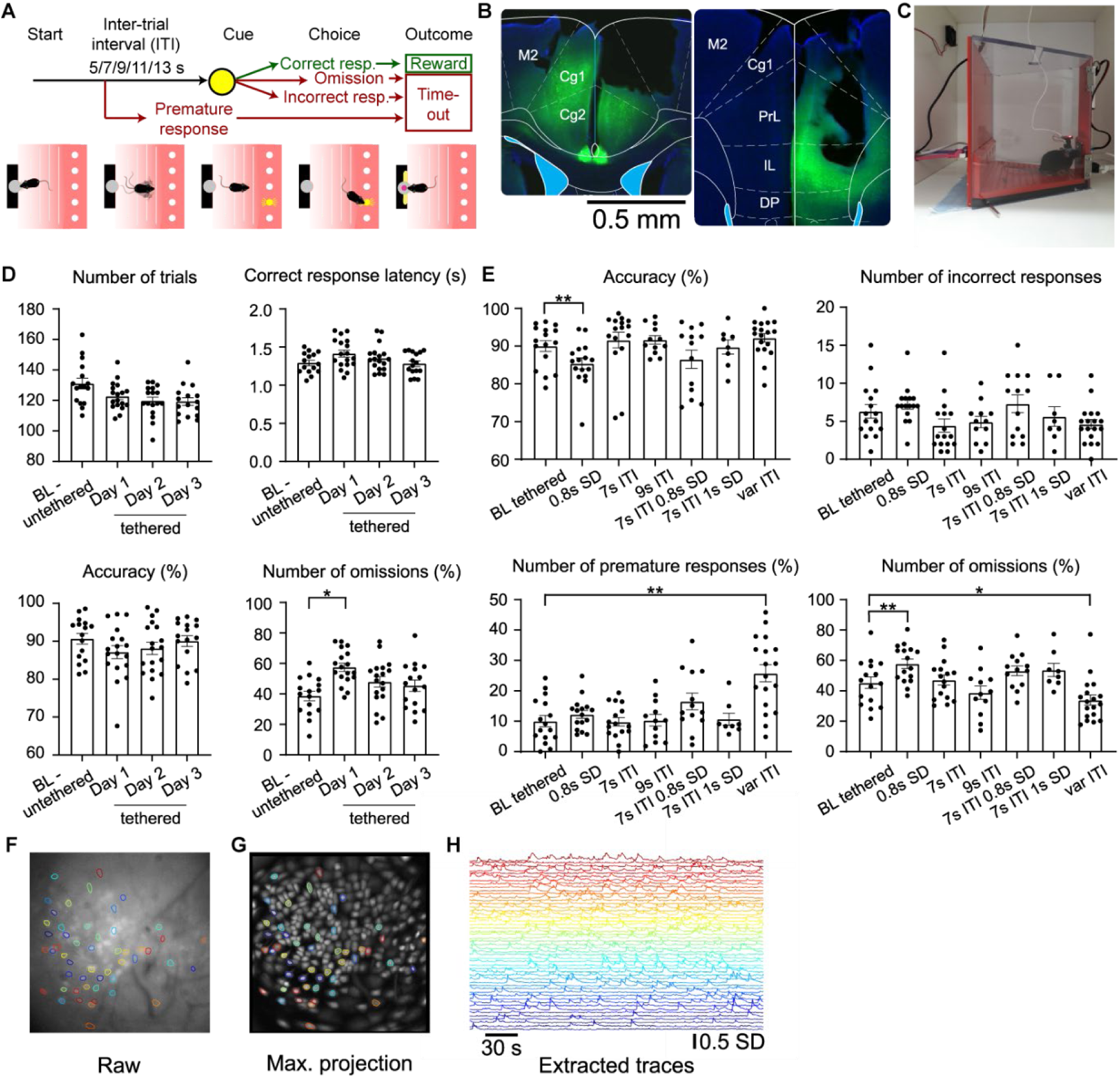
Behavioural performance with simultaneous miniscope recording. (**A**) Structure of an individual trial of the 5-CSRTT (see Methods for description). (**B**) Selective transfection of the ACC (comprised of regions Cg1 and Cg2; left, AP 1 mm) and the ventral mPFC (at the border between the regions PrL and IL; right, AP 2 mm) with GCaMP6m expressed in excitatory cells; black gap in the right hemispheres indicates GRIN lens location. (**D**) Measures of task engagement and performance in the 5-CSRTT (as indicated above panels) during training sessions without miniscope or dummy (baseline, BL-untethered) and during the first three sessions performed with tethered miniscope (Day 1-3). Dots indicate individual animals, bars show mean ± s.e.m.. Asterisks represent Dunnett pairwise post-hoc test comparing tethered days against baseline after significant effect of day in a one-way ANOVA. (**E**) Key performance indicators of the 5-CSRTT (as indicated above panels) measuring attention (accuracy, incorrect responses), impulse control (premature responses) and task engagement (omissions) for the baseline protocol and six challenge conditions during which miniscope recordings were conducted. Same display of mean ± s.e.m. and statistics (comparison of each challenge against baseline) as in (D). (**F**) Example of a 400 μm x 400 μm raw image obtained from ACC of an individual mouse during the 5-CSRTT with a miniscope, with exemplary identified active neurons encircled in different colours corresponding to traces shown in (H). (**G**) Similar display as in (F), but overlay of fields of view as maximum projection from 12 animals as imaged in the same 5-CSRTT protocol in ACC (circles indicate the same neurons as in (F). (**H**) Example of z-scored calcium activity traces over 10 min measured with GCaMP6m in the FOV shown in (F) from exemplary individual active neurons.

The recording of neural activity with miniscopes during operant tasks constitutes a challenge due to the relatively large and protruding form factor of such microscopes and a disadvantageous design of many operant box systems with deep and low recesses constituting the reward receptacle and poke holes. To enable miniscope recordings during the 5-CSRTT, we designed a custom-made operant box system with shallow and elevated poke-holes that reside in a protruding inner wall-layer (Figure 1C) [19]. This allowed mice to conduct the task with little disturbance by the mounted and tethered miniscope (UCLA model v3 or v4; Supplementary Video 1), as was confirmed by a lack of changes of achieved trial numbers, response latency, attentional accuracy (number of correct responses/(number of correct and incorrect response), and omissions (number of trials with omitted responses relative to total number of trials, %) beyond the first day of tethered training (Figure 1D). With repeated tethered training, animals performed well over 100 trials with less than 50% omissions on average, providing sufficient numbers of active responses for further analysis (Figure 1D).

In order to maximally engage attentional and impulse control – and to obtain sufficient numbers of incorrect and premature responses per session for later analysis - we performed six behavioural challenges with simultaneous miniscope recordings; this included a further shortening of the stimulus duration (SD) from 2 s at baseline to 0.8 or 1.0 s in challenge conditions, and/or an extension of the waiting time (inter-trial interval, ITI) before stimulus presentation from 5 s at baseline to fixed durations of 7 or 9 s or to variable lengths (7, 9, 11, or 13 s randomly at equal distribution, varITI). As expected, attentional performance, as indicated by accuracy, was lower with decreased stimulus duration (0.8 s SD challenge), which, however, also increased omissions, making it less suitable for analysis (Figure 1E). Overall, the varITI challenge appeared to produce the most suitable dataset for further physiological analysis, given that the relative number of premature responses was increased (*P* < 0.0001; Dunnet’s post-hoc test after significant main effect of challenge in mixed-effects ANOVA, *N* = 18) and the relative number of omissions was decreased (*P* < 0.001) compared to the baseline protocol, whereas incorrect responses were still present at a level comparable to the other test conditions (Figure 1E). We obtained stable recordings over 30 min sessions, yielding 10-72 cells per field of view (FOV) in the ACC and 14-72 cells/FOV in the mPFC (see Methods for details on trace extraction; Figure 1F-H).

### Individual ACC neurons have time-locked activity peaks around correct and premature responses

We first investigated qualitatively, if identified neurons display activity that is related to any of the four behavioural response options. Therefore, the calcium signal traces of each neuron were extracted from 4 s before until 7 s after each behavioural choice-poke event (note that, for omissions, the end of the stimulus presentation, was used as reference time point of choice for all time-locked analysis). Such individual episodic traces split into two populations of traces from trials with either *even* or *odd* order number; the averages of traces from *even* trials were then plotted in vertical order according to the peak latency of the averages of the corresponding *odd* trials (Figure 2A-B; Supplementary Figure 1). This indicated that ACC neurons often showed activity patterns that were time-locked to correct and premature responses. In support of this conclusion, Pearson correlations between the averages of *odd* and *even* trials indicated a high reproducibility of time-locked activity around *correct* and *premature* responses, with particularly high correlations (> 0.8) around and after the time of choice, in ACC and mPFC (Figure 2A-B, bottom; Supplementary Figure 1). Such correlated patterns were largely absent for incorrect responses and omissions. This constitutes a first indication that neural activity in both structures represents aspects of choices related to high attention (correct responses) and impulsivity (premature responses).

**Figure 2.**
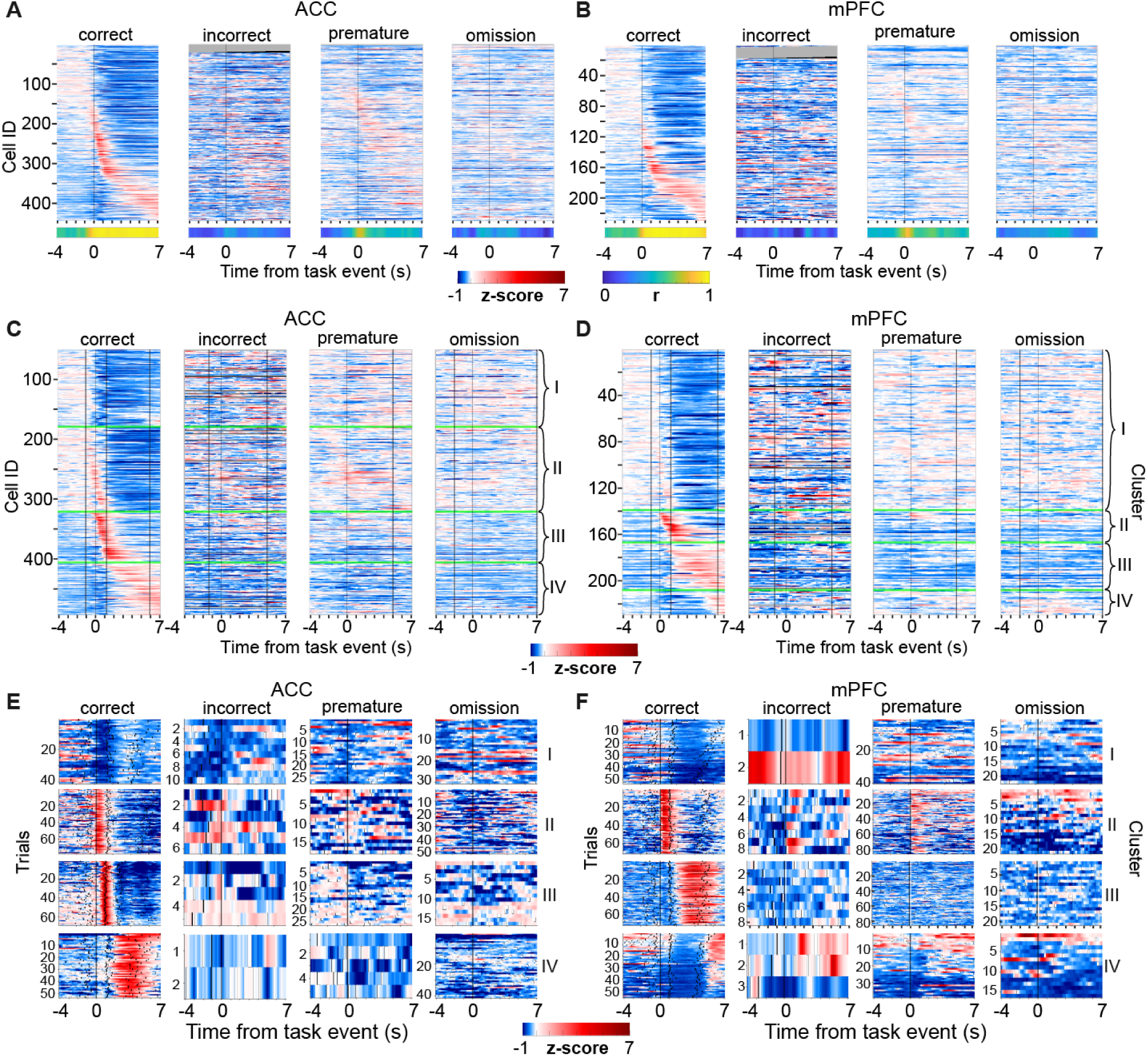
Event-locked activity of individual neurons in the variable ITI challenge. (**A**) *Top:* Average z-scored calcium activity in individual ACC neurons time-locked to the onset of the behavioural event stated above each sub-panel, shown for -4 - +7 s around the event. For cross-validation, averaging was done across the even trials only and the cells sorted according to the average peak latency across the odd trials (see also Supplementary Figure 1). *Bottom:* Pearson correlations between the averaged z-scored activity of the odd and even trials at each time point. Note that temporal order of peaks is maintained for correct and premature responses with resulting high correlations, but not for incorrect choices and omissions. *N* = 12 animals and *443* cells. (**B**) Same display and analysis as in (A) but for all neurons recorded in mPFC. *N* = 6 animals and *229* cells. (**C**) Same data as in (A) but clustered into four clusters (I-IV, indicated on the right) according to their activity from -4 - 7 s around correct responses. Equivalent to (A), clustering was done based on the average of *odd* trials, only, whereas the plot shows the average of the corresponding *even* trials. Activity around incorrect, premature and omitted responses for the same cells is shown in the same order as was determined according to the clustering around correct responses resulting in a lack of emerging temporal patterns. (**D**) Same analysis as in (C) but for cells in mPFC. Grey lines in (A-D) indicate cells for which a response cannot be shown because the mouse made no incorrect response. (**E**) Z-scored single-trial calcium activity of exemplary individual cells from each cluster (I-IV, indicated on the right). The timepoint of cue presentation (for correct and incorrect responses only), reward receptacle entry and exit (only correct responses) are shown by the white short vertical lines. The consistent white line represents the timepoint of the choice. (**F**) Same as in (E), but for cells from the mPFC.

Notably, correct and incorrect responses (in contrast to the other two event types) involve a similar global sensory stimulation (one poke-hole illuminated at the time of responding) and the same motor output (poking), suggesting that the time-locked activity seen only for correct responses, does likely not reflect sensory or motor aspects. However, we cannot fully exclude the possibility that deviations in the *local* stimulus - illuminated vs. non-illuminated hole into which the mouse pokes - may partially account for such differences.

To further investigate the temporal relationship of neural activity, we k-means-clustered the cells [20] into four clusters according to their average activity in *odd* correct response trials, sorted neurons within each cluster according to time of peak-activity, and then displayed the corresponding average of *even* trials (Figure 2C, D). Qualitatively, this resulted in three clusters with relatively clear activity peaks either before, during, or after correct responses, respectively, in addition to a fourth cluster with increased activity during reward collection only, in ACC (Figure 2C). In contrast, in mPFC a cluster with a well-defined peak at the time of the correct choices was lacking, and the emerging clusters showed activity either before or after the response (Figure 2D). To assess the response-specificity of these temporal patterns, we conducted the same temporal alignment and averaging for the other three response options but sorted the cells according to their order number obtained for clustering by correct responses. For all three response types, this resulted in the loss of clear temporal response patterns, indicating that the temporal relationship cells displayed for correct responses, were largely specific for this one response type (Figure 2C-D). Finally, when plotting the activity in individual trials of one randomly selected neuron for each cluster, the trial-to-trial reliability of activity as time-locked to correct responses was qualitatively confirmed (Figure 2E-F).

### Distinct choices are represented by ACC population activity

While the analyses described above confirm that individual ACC and mPFC neurons are modulated by ongoing attention- and impulsivity-related choices or actions, a comprehensive and multi-variate encoding of behaviour is expected only at the level of populations of multiple neurons. To evaluate such behavioural representations in ACC, we performed a decoding analysis, training linear support vector machine (SVM) classifiers to predict the type of behavioural event performed in any given trial based on the population activity, at different time points around the choice event, and during the first 4 s of the preceding ITI (starting with the end of the time-out or of reward consumption). We first focused on binary discrimination between correct and either omitted or premature responses, given that incorrect responses occurred in insufficient numbers for this analysis, in the varITI-challenge in most mice (Figure 1E; Supplementary Table 1). To estimate significant predictions, we compared the resulting accuracies with accuracies obtained from classifiers that were trained on data with randomly shuffled labels (paired *t*-tests at each time point with Benjamini-Hochberg correction for multiple comparisons).

No appreciable prediction of correct responses (vs. omissions or vs. premature responses) was possible during the ITI, indicating that ACC activity did not reflect, if a mouse *was going to* act in a goal-directed, attentive fashion or to avoid task engagement or to act impulsively in an upcoming trial (Figure 3A, left). Although, there was a significant decoding of correct vs. omitted or premature responses at low accuracy of around 60% (vs. 50% chance level), already from at least 4 s before the choice poke onwards, average decoding accuracies only started rising around cue-onset and reached their maximum of >90% only approx. 600 ms *after* the choice-poke. They remained at >90% throughout the time of reward consumption (Figure 3A, right). This indicates that the pre-cue and pre-choice representations were very minor compared to the same representation around and after the choice. These results appear inconsistent with the notion that the primary driver of variance in ACC activity are slowly varying cognitive states of attention or impulsivity, but rather that ACC representations seems to be tightly tied to actions and outcomes. In mPFC, decoding accuracies had a similar temporal trajectory (Figure 3B).

**Figure 3.**
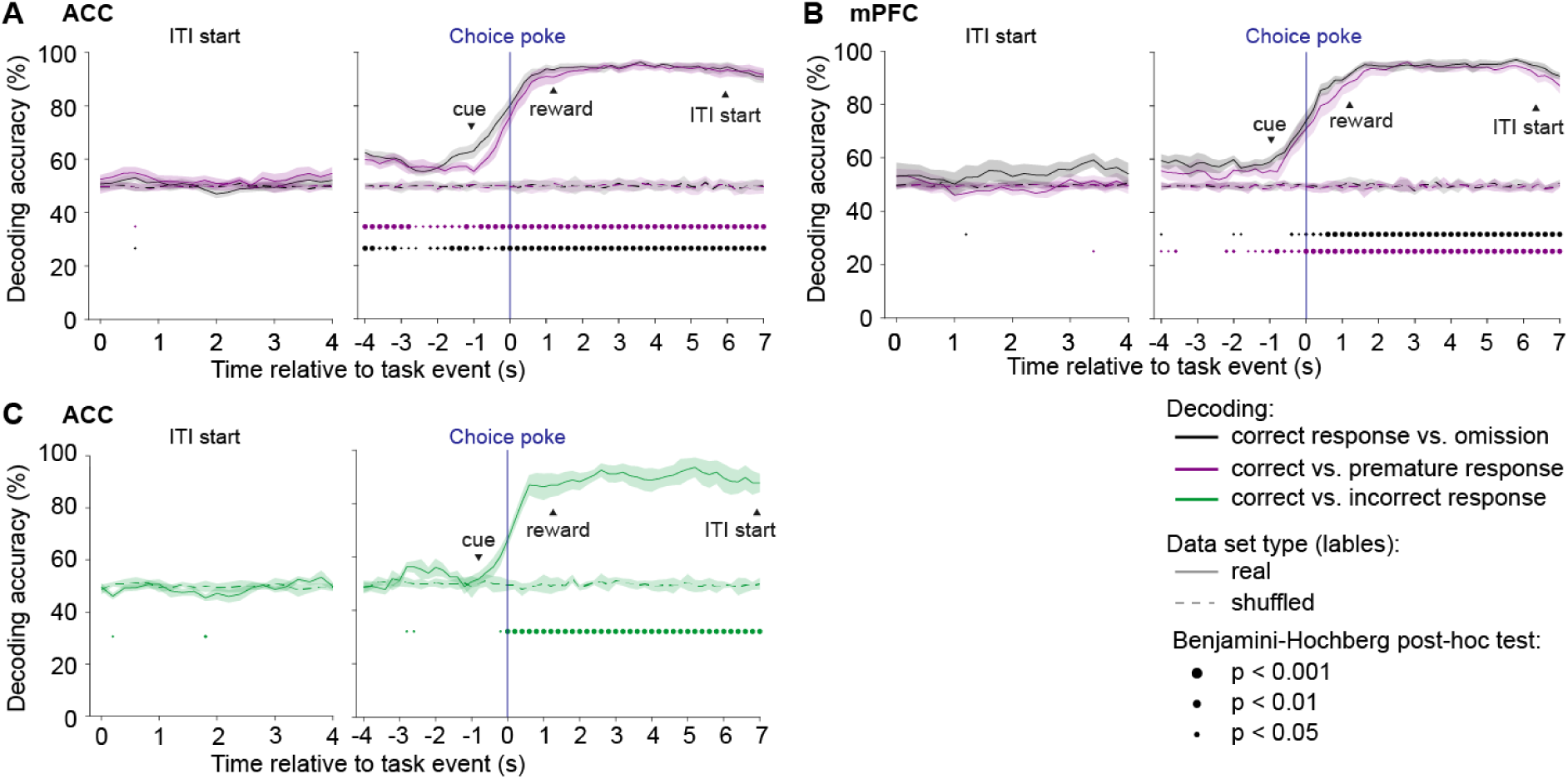
Decoding of behavioural choice from population activity in ACC and mPFC. (**A-B**) Cross-validated decoding accuracies derived from binary classification using linear SVMs calculated from z-scored amplitude values at each 200 ms time bin and predicting behavioural events from population activity in ACC (A; *N* = 11 mice; one mouse not analyzable due to low omission rate) or mPFC (B; *N* = 6 mice) in individual varITI sessions; solid lines and shading represent averages across animals ± s.e.m., respectively. Decoding accuracies were first averaged across 100 classifiers calculated on data from each session, and then across sessions (i.e. animals). Dashed lines indicate results from the same analysis but performed on control data obtained by random shuffling of event-labels relative to neural activity data; dots at the bottom indicate a significant difference in the pairwise comparisons between those two accuracy values at each time point (*t*-test with Benjamini-Hochberg correction for multiple comparisons). Binary classification was done differentiating correct responses against omissions (black) or against premature responses (magenta). Chance level is 50%. (**C**) Same analysis as in (A) but classifying *correct* vs. *incorrect* responses by using sessions from across all challenge protocols, if more than 5 incorrect responses were made (*N* = 6). mPFC was not analysed because only 3 sessions had the sufficient number of incorrect responses. See Supplementary Table 1.

In the pairwise discriminations described above, a confound by non-choice-related aspects such as presence of the cue (*correct* vs. *premature*) or the motor response (*correct* vs. *omission*) cannot be ruled out. Only *correct* and *incorrect* responses are sufficiently similar in most parameters and differ mainly in the choice *per se*. To enable a cross-validated decoding analysis involving *incorrect* responses, we gathered sessions from all six behavioural challenge conditions given they had a sufficient number of incorrect responses (≥ 6). Using such data, we found that – in contrast to premature and omitted responses - incorrect choices could be distinguished significantly from correct ones only from the time of the choice-poke onwards (approximately 1 s after cue-onset), indicating a lack of a consistent signature of both preparatory attention and of sensory stimuli in the overall ACC activity (Figure 3C). As seen with the other decoding attempts, high and saturating average prediction accuracies (>80%) could only be reached from around 600 ms after the choice-poke and were maintained throughout reward consumption or its omission (Figure 3C). Overall, the temporal distribution of decoding accuracies is more aligned with the notion that ACC represents ongoing action and its (expected or actual) consequence rather than controlling levels of sustained attention or impulsivity on a trial-by-trial basis.

### ACC neuron populations encode spatial aspects of ongoing action

The observation that decoding of response types available in the 5-CSRTT was only possible with high accuracy from the choice-poke onwards, suggests that ACC and mPFC may represent selected actions rather than high-level cognitive states like attention and impulsivity. To further scrutinize this hypothesis, we investigated whether these neurons encode a more fine-grained representation of current action by analysing the responses to each individual choice poke-hole. We aligned the average activity of each neuron in *even* trials to the time of correct choice for each hole individually and sorted the neurons first according to the hole (1-5) which evoked the strongest response during the poke (±1 s) into five groups, and then sorted by peak latency of the average of *odd* trials within each group. A reproducible pattern emerged for *even* trials that correlated strongly to that of *odd* trials from around the time of choice onwards, in ACC and mPFC (Figure 4A-B, bottom), and which appeared the more dissimilar between poke-holes around the time of poking, the further the holes were apart from each other (Figure 4A-B, top).

**Figure 4.**
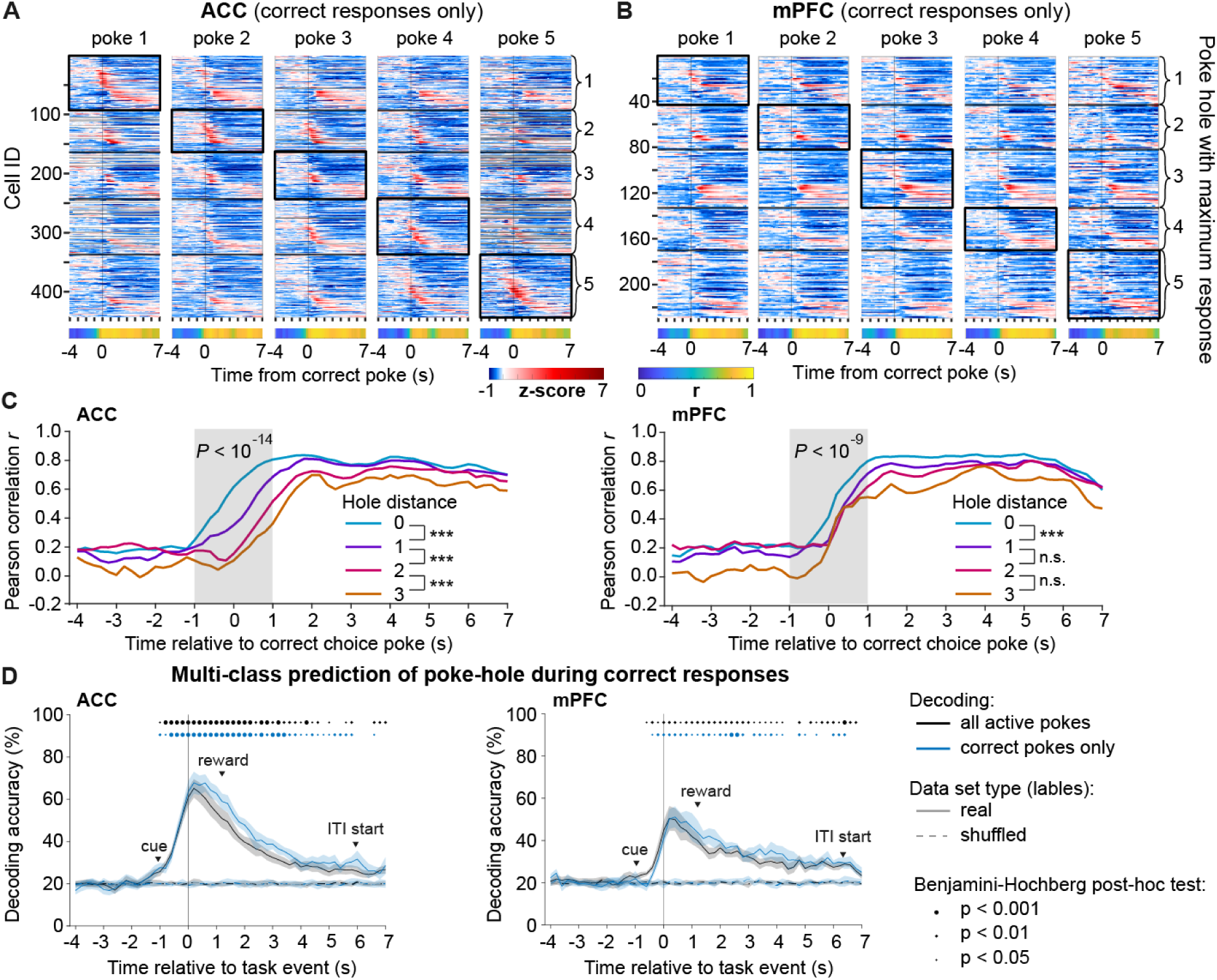
Activity in the ACC represents spatial action selection. (**A-B**) Same data as in Figure 2A-B (correct responses), but separated by poke-hole and arranged into five groups (indicated on the right, separated by black lines) based on the hole for which the response of a neuron had the maximum AUC in the period ±1s around the poke (indicated by a black rectangle around the cluster). *Top:* Average z-scored calcium activity in individual ACC (A) or mPFC (B) neurons time-locked to the correct choice-poke, shown for -4 - +7 s around the event. For cross-validation, averaging was done across the *even* trials only and the cells were sorted according to the average peak latency across the *odd* trials of poke 1. Grey lines indicate sessions in which the given hole was not poked into. *Bottom:* Pearson correlations between the averaged z-scored activity of the *odd* and *even* trials at each time point. Note that the qualitative similarity to the pattern of a given poke gets reduced the further away the poke-hole is, especially around the time of poking. Supplementary Figure 2A-B shows the same data without prior sorting into clusters. (**C**) Based on the data shown in (A-B), Pearson correlations between response patterns of pairs of poke holes, coded in colour according to the distance between the holes; correlation values for hole-combinations with the same distance (e.g. 1-4 and 2-5 for distance 3) were averaged; see Supplementary Figure 2C for the individual correlation values of each hole-pair. A repeated-measures ANOVA was calculated across the population of 11 observations in the time period ±1 s around the poke (grey bar; *P*-value for main effect of hole-distance indicated at the top); results from paired Sidak post-hoc tests are indicated for adjacent hole distances in the colour-legend of each sub-panel. *** *P* < 0.001; n.s., *P* > 0.5. (**D**) Accuracies of the decoding of the identity of the poke-hole, either for all active responses (correct, incorrect, premature; black) or for correct responses only (blue) aligned to the time of poking (0, vertical line); average latency to cue onset (response latency), reward-poke entry (reward latency) and start of the next ITI (reward consumption time, indicted only for correct trials) are indicated by arrowheads. Decoding accuracies were first averaged across 100 classifiers calculated on data from each session, and then across sessions (i.e. animals). Dashed lines indicate results from the same analysis but performed on control data obtained by random shuffling of event-labels relative to neural activity data; dots at the top indicate significant pairwise comparisons between those two accuracy values (*t*-test after correcting for multiple comparisons across time bins). Shaded area, s.e.m. across mice.

To assess this quantitatively, we calculated Pearson correlations between such population activity patterns for all pairs of poke-holes and averaged them across hole-pairs with the same distance (Figure 4C; Supplementary Figure 2). Indeed, within approximately ±1 s around the choice-poke, correlations were the higher the closer the holes of the pair were to each other. After the choice, in contrast, correlations were consistently high, irrespective of distance. This suggested that ACC activity displays a certain similarity related to spatial proximity of poke holes before the poke, while being dominated by non-spatial aspects of the choice after it is made (Figure 4A-C).

To further investigate this early spatial selectivity, we trained multi-class SVM classifiers to decode the identity of the poke-hole. Surprisingly, this identity could be predicted from about 1 s before the poke (just after cue-onset) onwards, reaching an average peak accuracy of close to 70% in ACC and close to 55% in mPFC (against a chance level of 20%) approximately 200 ms after the poke. In contrast to the representation of event-type (Figure 3A-C), average accuracy decreased again immediately after the poke, suggesting that the representation of the precise action fades after its occurrence (Figure 4D). Interestingly, the decoding accuracy was virtually identical, when performing the same analysis on pokes of all three active response types (correct, incorrect, premature) *combined*, suggesting the existence of a representation of action that is independent from the representation of response-type and outcome.

### Independent encoding of action and outcome in ACC

The hypothesis of an independent encoding of motor action (poke hole identity) and choice or outcome (event-type) entails the question to what extent each factor determines neural activity in the ACC, at every time point. To answer this question, we trained linear regression models predicting the activity of each neuron using predictors coding the active poking (vs. omissions), rewarded choice (correct responses vs. absence of reward due to erroneous choices), and poke-hole identity (spatial location; encoded by four predictors, see predictor matrix in Figure 5A). We ran a separate regression for each time-point in the aligned activity and calculated the cross-validated coefficient of partial determination (CPD) for each predictor at each time-point, which quantifies the share of the variation in the neural activity that is uniquely explained by that predictor. To estimate statistical significance for a given predictor at a given timepoint, the distribution of CPD values across subjects for that predictor and timepoint was compared to 0% [21].

**Figure 5.**
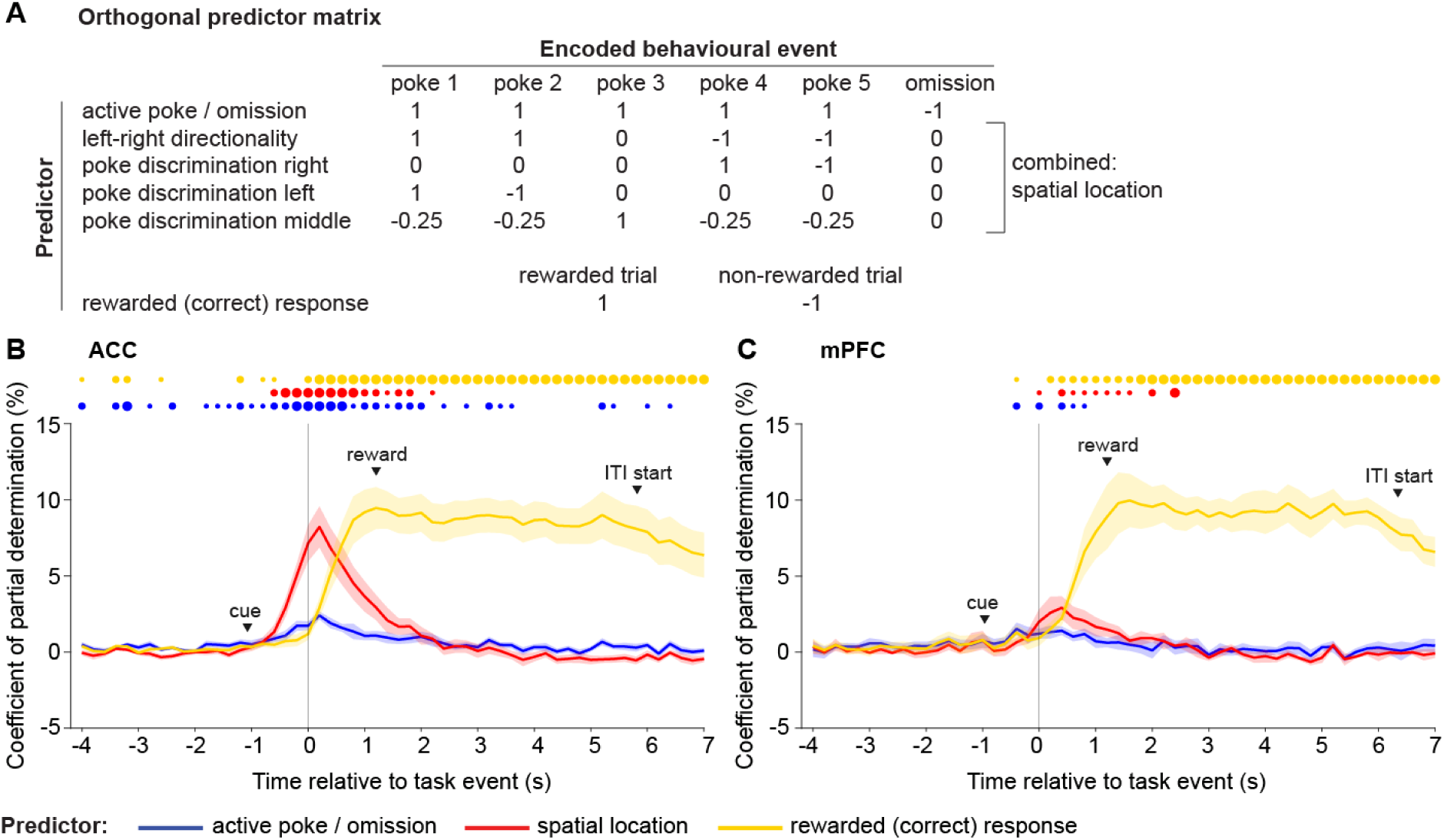
Encoding of poking action and reward in the population activity of the ACC and mPFC. (**A**) Orthogonal predictor matrix designed to indicate the representation of the poke (1: poke in either hole, 0: omission), the poke directionality (1: poke in left holes 1 or 2; - 1: poke in right holes 4 or 5; 0: poke in middle hole 3 or omission), right poke discrimination (1: poke in hole 4; -1: poke in hole 5; 0: poke in holes 1,2 and 3 and omission), left poke discrimination (1: poke in hole 1; -1: poke in hole 2; 0: poke in holes 3,4 and 5 and omission), middle poke discrimination (1: poke in hole 3; -0.25: poke in holes 1,2,4,5; 0: omission) and reward (1: rewarded; 0: not rewarded). The four predictors representing the spatial location of the poke hole have been removed at once from the regression model in order to quantify the encoding of motor action. (**B**-**C**) Coefficient of partial determination (CPD) averaged across cells recorded in ACC (B, *N* = 11 mice) and mPFC (C; *N* = 6 mice). Time bins where CPDs for a given event were significantly higher than zero after cross-validated linear regression are indicated with a dot at the top of each panel, colour-coded for the respective predictor (one sample *t*-test with Benjamini-Hochberg post-hoc correction). CPDs were determined for each event-type by subtracting the sum squared errors of the full linear regression model (incorporating every event type as predictor) from the sum squared error of the reduced regression model where one predictor (corresponding to the event type for which the population activity should be explained) was removed.

From the average onset of cue presentation (1 s before the poke), matching the time course of the decoding analysis (Figure 4D), the spatial identity of the poke-hole started to gain ever more influence over ACC-activity, and dominated it compared to the other predictors from approximately 600 ms before until 400 ms after the choice poke (Figure 5B). This effect was mostly driven by an encoding of the left (hole 1-2) vs. the right (hole 4-5) side of the 5-choice wall, although most other tested predictors of the selected action (poke discrimination left, right, and middle) and of active responding in general (vs. omission) displayed significant CPDs during and after the time of poking, as well (Supplementary Fig. 3). From 600 ms after the poke onwards, however, the factor of rewarded (correct vs. erroneous) response had the single strongest influence on neural activity out of the tested predictors (Figure 5B). This overall pattern suggests, that AAC *simultaneously* encodes fine-grained selected action and a high-level representation of both active and correct responding from the time of choice-poke onwards for almost 2 s, but action representation dominates around the time of execution whereas high-level representation dominates subsequently (Figure 5B). Given that the influence of outcome starts rising immediately after the choice poke - and hence more than a second before actual reward consumption - this activity reflects *expected* (vs. omitted) reward at this early stage, but it might later on reflect actual outcome, as previously shown [11]. mPFC neurons, in contrast, also encoded outcome (from 200 ms after the poke) but lacked a consistent encoding of motor action, especially before the poke (Figure 5C).

### Devaluation and omission of reward alter representation of choices

The prominent representation of *correct* – and, hence, *rewarded* – responses in ACC, particularly during and after the response, which is also in line with earlier studies [9,10], suggests that these activities may represent the expectation of reward rather than elevated, preparatory attention. This is also supported by the fact that incorrect and correct responses cannot be discriminated from population activity *before* the choice-poke is made (Figure 3C). To assess this conclusion further, we conducted four further 5-CSRTT experiments where the value or expectancy of the reward was altered, using the combined 0.8s/7s ITI challenge (to obtain more incorrect responses, Figure 1E), at the end of the sequence of tests: first we recorded a baseline session with normal food-deprivation and reward delivery. In a second session, the reward was *devalued* by pre-feeding (by providing 6 g of food overnight and 2 ml of milk reward 1 h before session start). In the third and fourth recording sessions the food-deprivation (i.e. value of reward) was normal, but the delivery of reward was omitted (*extinction* 1 and 2). Normal training sessions in the baseline protocol were conducted after the first two sessions, but not between the extinction sessions.

At the behavioural level, both devaluation and extinction caused a significant decrease of the number of correct responses, driven by an increase in omissions, whereas accuracy remained relatively stable (*P* < 0.05, Dunnett’s post-hoc test, performed after significant main effect of condition in RM-ANOVA; *N* = 15; Figure 6A-C). During devaluation, also active *erroneous* (incorrect and premature) responses decreased, and reward latency increased, in line with a reduced motivation (Figure 6D; Supplementary Figure 4); such effects appeared qualitatively also during extinction sessions, but mostly without reaching significance. Response latencies were not significantly altered indicating unperturbed responsiveness in any of the conditions (Supplementary Figure 4).

**Figure 6.**
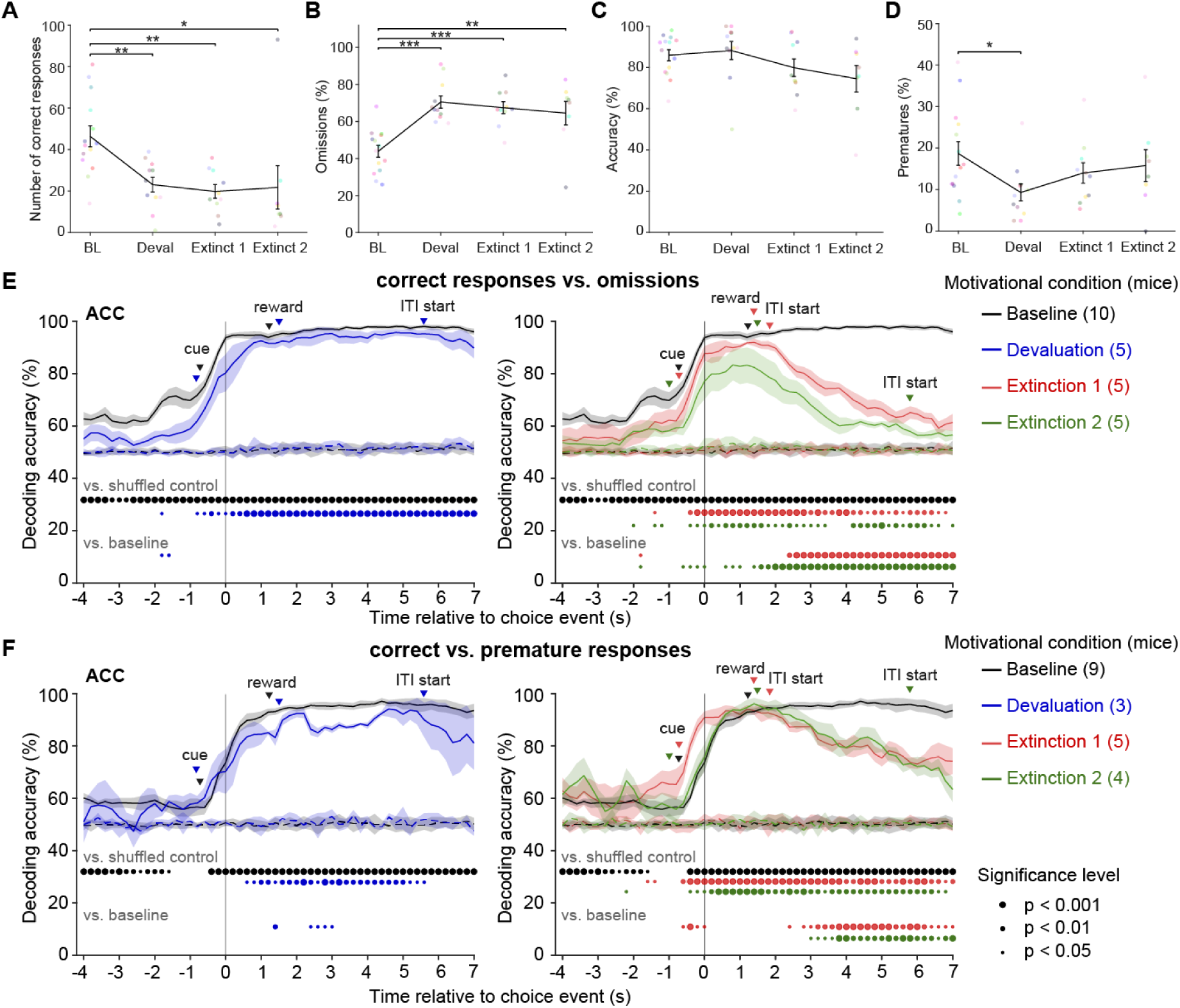
Decoding of behavioural choice from population activity in ACC during reward devaluation and extinction experiments. (**A-D**) Measures of task engagement and performance in the 5-CSRTT (as indicated on y-axes) during training sessions in the 0.8s-SD/7s-ITI combined challenge at normal reward conditions (baseline, BL), after devaluation of reward (Deval), or with omission of reward (extinction; Extinct 1/2), as indicated on x-axes with tethered miniscope. Dots indicate individual animals coded by colour, bars show mean ± s.e.m.. Asterisks represent Dunnett pairwise post-hoc test comparing BL condition against the other conditions after RM-ANOVA. * *P* < 0.05; ** *P* < 0.01; *** *P* < 0.001. See Supplementary Figure 4 for further behavioural measures. (**E-F**) Cross-validated decoding accuracies derived from binary classification using linear SVMs calculated at each 200 ms time bin and predicting correct vs omitted (E) or vs. premature (F) responses from population activity in ACC during test sessions in the 0.8s-SD/7s-ITI combined challenge with normal reward conditions (baseline, black), after devaluation of reward (blue) or with omission of reward on two consecutive test sessions (extinction 1, red; extinction 2, orange). Whereas 10 mice participated in these test sessions, actual *N*-numbers for the analyses (stated in figure legends on the right and in Supplementary Table 1) vary mainly due to mice that did not perform sufficient numbers of correct or premature responses, in rare cases also due to technical failures. Solid lines and shading represent averages across animals ± s.e.m., respectively. Decoding accuracies were first averaged across 100 classifiers calculated on data from each session, and then across sessions (i.e. animals). Dashed lines indicate results from the same analysis but performed on control data obtained by random shuffling of event-labels relative to neural activity data. Dots below the traces of each panel indicate time bin and alignment (coded by colour) where decoding accuracies from these two classifiers differed significantly (paired *t*-test with Benjamini-Hochberg correction; coded by colour for each reward condition). Dots at the bottom represent Dunn-Sidak post-hoc tests comparing accuracy values from devaluation or extinction sessions (coded by colour) to those from the baseline condition (conducted after significant effect of reward condition in the repeated-measures ANOVA). Significance level is encoded by dot size. Triangles indicate average latency to cue-presentation, reward receptacle entry and the start of the next ITI (the latter was often close after reward receptacle entry in extinction conditions as no reward was provided). Chance level is 50%. Decoding of incorrect responses could not be performed due to their small numbers in most of the respective sessions.

To evaluate if the change of the value or contingency of the reward affected the representation of rewarded responses, we repeated the time-resolved binary decoding of correct vs. premature or omitted responses, as conducted previously for the varITI-challenge (Figure 3), in each of the four conditions. Whereas, under normal reward conditions, the trajectories looked like those found before, with increasing representation of the choice and its outcome with the cue-onset, somewhat altered decoding accuracies were found in the other conditions (Figure 6E-F). Generally, decoding accuracy was lower in the conditions with altered reward value or occurrence, as indicated by comparisons to the accuracy achieved by classifiers trained with shuffled control datasets. Significant decoding of correct responses was hardly possible *before* they actually occurred, possibly reflecting a certain lack of representation of preparatory attentional or impulse control required for such responses (Figure 6E-F). When comparing the accuracy achieved under baseline condition with that achieved under reward devaluation (RM-ANOVA with pairwise Dunnett post-hoc tests), these two conditions rarely differed, suggesting that the value of the reward has relatively little influence on the discriminability of representations of rewarded vs. non-rewarded choices (Figure 6E-F). In contrast, during both *extinction sessions*, the discrimination of correct responses vs. omissions and, to a lesser extent, vs. premature responses, deteriorated from about 2 s after the choice poke onwards and differed significantly from the accuracy achieved in the baseline condition, in line with the lack of a reward (Figure 6E-F). This implies that beyond this time point, ACC largely encodes the *actual* outcome, which is no longer distinct between the choice options in the extinction sessions. Similarly, in the *second* extinction session, accuracy for decoding correct responses vs. omissions differed already from 600 ms *before* the choice poke onwards and on most subsequent time points (Figure 6E). This supports the idea that ACC partly represents *expected* outcome, which has been learned to be identical between these two choice options by the time of the second extinction session.

To further elucidate the hypothesis that ACC encodes *expected* and, subsequently, *actual* outcome, we performed an encoding analysis, as previously done for the varITI-challenge (Figure 4), for the data obtained from these four conditions. Across time intervals, we compared CPD values to the control value of 0% for all conditions (*t*-test with Benjamini-Hochberg correction) and we compared the devaluation and extinction conditions to baseline (RM-ANOVA followed by Sidak post-hoc test; Figure 7A). As expected, *extinction* – especially when repeated – led to a reduction of the representation of reward. However, also *devaluation* ensued a strong decrease of reward representation compared to baseline, suggesting that reward determines ACC activity the stronger the higher its value is (a result that the prior decoding analysis could not reveal, see Figure 6E-F). Strikingly, both devaluation and extinction also led to a virtual loss of the spatial representation of the poke-hole: in all three conditions, CPD values for the combined spatial predictors were significantly lower than those in the baseline condition around the time of poking and, in contrast to the baseline, rarely exceeded 0% (Figure 7A). This suggests that the representation of specific actions in ACC depend on their expected (reward) value.

**Figure 7.**
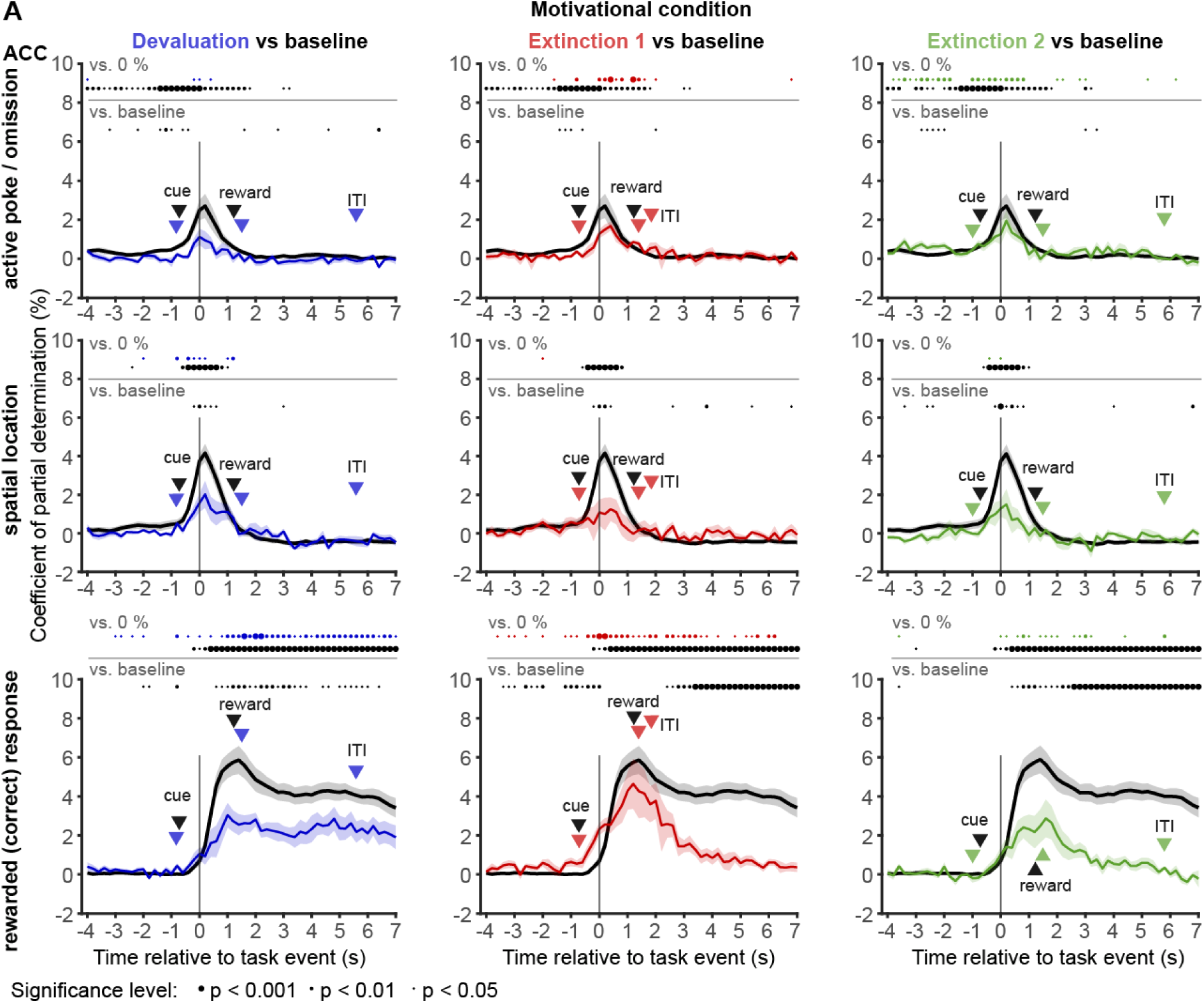
Encoding of reward and action in ACC depends on the relative value and presence of reward. (**A**) Coefficient of partial determination (CPD) averaged across cells recorded in ACC in the four different reward-related conditions encoded by colour (number of animals): black, baseline (10); blue, devaluation (6); red, extinction session 1 (6); green, extinction session 2 (6). As in Figure 5, CPDs were determined for each predictor, stated on the left (rows) by subtracting the sum squared errors of the full linear regression model (incorporating every event type as predictor) from the sum squared error of the reduced regression model where one predictor was removed. To evaluate the encoding of spatial location as such, all four predictors reflecting spatial location were combined by removing them at once (see Figure 5A). Time bins where CPDs for a given event were significantly higher than zero after cross-validated linear regression are indicated with a dot at the top of each panel, colour-coded for the respective reward condition (one sample *t*-test with Benjamini-Hochberg post-hoc correction). Dots below represent Sidak tests comparing CPD values from devaluation or extinction sessions (coded by colour) to those from the baseline condition (conducted after significant effect of reward condition or time-reward interaction in RM-ANOVA). Significance level is encoded by dot size. Triangles indicate average latency to cue-presentation, reward receptacle entry and the start of the next ITI (the latter was often close after reward receptacle entry in extinction conditions as no reward was provided). To ensure, that CPDs in the baseline session are not simply higher because of a larger number of trials or responses used for their calculation, the number of baseline-trials was down-sampled to roughly match those of the reward condition it is compared to in each panel for each poke-hole and response type.

## Discussion

Using miniscope recordings during the 5-CSRTT in mice, we here demonstrated that excitatory neurons of the posterior ACC represent the choices available in this complex task, both at the high level of response options that depend on *cognitive state* and at the low level of the selected *action* (poke-hole identity). We found that coding at both levels happens simultaneously, but that ACC activity is dominated by the selected action (graded spatial representation of poke-hole location) during the execution of the choice poke, and by outcome expectation from about 400 ms after the choice, followed by encoding of actual outcome, once obtained. Whereas action representation faded, the high-level representation of the chosen option (or its consequence) remained stable for at least 7 s after the choice, but deteriorated much faster when reward was omitted during extinction sessions. When value or occurrence of reward were changed, the share of cells selectively *activated* by reward and of cells selectively *inhibited* during omissions – i.e., the share of cells representing those options whose relative utility changed – increased. This suggests that ACC represents action-outcome contingency rather than outcome itself. Likewise, the representation of rewarded (correct) responses, but also of low-level spatial action parameters decreased when reward was either devalued or omitted. This underlines that motor and reward representations in ACC depend on the expectation and value of a choice’s outcome.

Importantly, significant preparatory network activity during the ITI and before the cue which could indicate a general level of high or low attention, impulsivity, or task engagement in a given trial was either not found (attention-related activity discriminating correct from incorrect choices) or was rather weak (discrimination between impulsive responses or omissions and correct responses). In fact, all conducted analyses assessing response-type coding suggest that choices are represented in ACC mainly *during* and *after* their time of occurrence, and that *rewarded* responses are represented most reliably, as seen in other studies [12,13].

How can these observations be reconciled with earlier reports of signatures of attention and impulsivity in ACC [9–11,22] and, most importantly, with the well-documented ability to modulate these cognitive states by manipulation of ACC neurons [5,6,8,23]? Firstly, with respect to pre-choice activity, the discrimination of correct responses from either premature or omitted responses *was* actually possible, albeit at low (∼60%) accuracy, already during at least 4 s before the choice poke (but not at the beginning of the ITI; Figure 3, 6). Furthermore, correct and premature responses were accompanied by time-locked activity of neurons in the same pre-choice time period (Figure 2). This indicates a representation of the state of impulse control that exists, albeit weakly, well before the choice, though not at the beginning of the ITI – in line with ramping activity before premature responses observed in rats [10]. In contrast, incorrect responses could only be distinguished from correct responses at time of occurrence (Figure 3) and were not accompanied by time-locked activity before (Figure 2). This suggests that the trial-to-trial variation of sustained attention is not represented in excitatory cells of posterior ACC. The divergence between both phenomena is in line with the fact, that impulsivity, but not accuracy, was modulated by chemogenetic manipulation of such neurons in ITI-challenges [5]. Since modulation of ACC parvalbumin-interneurons (which we did not record in this study) could alter both aspects of executive function [8], they could constitute a locus of the attentional component in this task [14]. Thirdly, chemogenetic or pharmacological modulation of ACC could exert its effect anatomically and temporally more globally, rather than controlling behaviour directly and in individual trials. The ACC might be acting through the demonstrated strong representation of obtained reward, errors, and action-outcome contingencies, to shape tendencies of attentional and impulse control in upcoming trials through other circuits, as shown for a subpopulation of ACC neurons already [6,7]. Fourthly, and alternatively to the previous scenario, ACC modulation may influence the occurrence of distinct response-types because of its role in action selection [21,24], as we suggested based on our [5,8] and other [6] previous chemogenetic and optogenetic [7] data before [8]. In this scenario, elevated activity of certain ACC pyramidal neurons triggers the response into a certain poke-hole (in line with the early encoding of action found in this study). This could be caused by the ACC due to reward expectation leading to strong excitatory AAC activity entailing a correct response. But it could also be caused erroneously by higher order inputs received by the ACC without the appropriate cue leading to incorrect or premature responses, accompanied by somewhat less effective, less time-locked ACC excitation. Activation of PV-interneurons [8] or direct partial inhibition of excitatory neurons in ACC [5] could increase the threshold needed for such activation that ultimately triggers the selected response, so that the strong activation triggering correct responses remains supra-threshold, whereas the weaker activity which could otherwise cause erroneous responses falls sub-threshold. Further physiological investigations are required to probe these mechanistic models.

Regarding mPFC, the number of mice we recorded in our study for comparative purposes is too small to draw general conclusions regarding the coding in this region, which has also been examined in other studies in the 5-CSRTT [10,14,26,27]. Nevertheless, the relatively minor differences between representations in ACC and mPFC we observed are consistent with an earlier finding of oscillatory coupling of both regions during choice events in the 5-CSRTT [22] and, more generally, with a model of widely distributed encoding of behaviour across neocortex [16]. Population activity in mPFC also represented choice options but less reliably and partly shifted to later time points, i.e., after the choice, and again with a bias to encoding rewarded choices more than non-rewarded ones. Also, a fine-grained encoding of spatial aspects of a selected action was virtually absent. This is in line with previous electrophysiological measurements in rats during this task, indicating that a majority of responsive mPFC cells respond after the choice, representing trial outcome, and cells show considerably stronger firing rate increases during correct than during premature responses [10].

In conclusion, our temporally resolved analysis suggests that ACC excitatory neurons represent a chosen action as it is made as well as its expected and actual outcome. Our results uncover parallel encoding of fine-grained spatial parameters of selected actions - in dependence on their outcome value - and of action-outcome contingency in ACC, and suggest that trial-by-trial encoding of high-level cognitive states before the choice is either minimal (for task engagement and impulsivity) or absent (for attention).

## Methods

### Animals

In total, 46 male C57BL/6J wildtype mice, were used for this study. Animals were group- or single-housed in Type II-Long individually ventilated cages (Greenline, Tecniplast, G), enriched with sawdust, sizzle-nest^TM^, and cardboard houses (Datesand, UK), and maintained at a 13 h light / 11 h dark cycle. Water was available *ad libitum*. All experiments were performed in accordance to the German Animal Rights Law (Tierschutzgesetz) 2013 and were approved by the Federal Ethical Review Committee (Regierungsprädsidium Tübingen) of Baden-Württemberg, Germany (licence number TV1469).

### 5-CSRTT training and testing with calcium imaging

Mice started training in the 5-CSRTT at 3-5 mo of age and were kept under food-restriction at 85-95% of their prior average free-feeding weight which was measured over 3 days immediately prior to the start of food restriction at the start of the behavioural training. Testing was conducted in operant chambers placed individually in melamine-MDF sound-insulated and ventilated outer boxes and fitted internally with an array of five nose-poke holes on one wall and a reward receptacle on the opposite wall. All six apertures could be illuminated to instruct the entry into them and were fitted with IR break-beams to detect entry and exist of the animal’s snout. All experiments were conducted in custom-made trapezoidal chambers based on the pyControl system [19,28] (https://pycontrol.readthedocs.io).

The 5CSRTT training protocol was similar to what we previously described [5]. In brief, after initiation of food-restriction, mice were accustomed to consume the reward (strawberry milk, Müllermilch^TM^, G) first in their home cage, and then in the operant box (2-3 exposures each). Subsequently, mice were trained in 2-13 sessions (30 min, once daily) of habituation training. In each trial, all holes of the 5-poke wall were illuminated for an unlimited time and the mouse could poke into any one of them to earn a 40 μl milk reward subsequently disposed from the illuminated receptacle. If mice attained at least 30 rewards each in two consecutive sessions or (in exceptional cases) had reached the 16^th^ session of habituation training, they were moved to the 5-CSRTT training, during which mice transitioned through five stages of increasing difficulty, based on reaching certain performance criteria in each stage, as described previously[5]. The difficulty was determined by the length of time the stimulus was presented (stimulus duration, SD) and the length of waiting time between the end of the previous trial and the stimulus presentation of the next trial (inter-trial-interval, ITI). In case a reward was collected on the previous trial, the ITI was initiated by the removal of the snout of the animal from the reward receptacle. In all 5-CSRTT protocols only one pseudo-randomly selected aperture of the 5-choice wall was lit up after the ITI, indicating that this hole needs to be poked into (*correct response*) in order to earn a 20 μl milk reward (Figure 1A). Trials were not rewarded but instead terminated immediately with a 5 s time-out period during which the house light was turned off, if the animals either poked into any hole during the ITI (*premature response*), poked into a non-illuminated hole (*incorrect response*) during the SD and limited-hold time (LH, until 2 s after SD), or failed to poke throughout the trial (*omission*). The relative numbers of such response types were used as performance indicators measuring premature responding [%premature = 100*(number of premature responses)/(number of trials)], sustained attention [accuracy = 100*(number of correct responses)/(number of correct and incorrect responses combined)], and lack of participation [%omissions = 100*(number of omissions)/(number of trials)]. A trial was considered to start at the beginning of the ITI, i.e. included premature responses. Additionally, the time required to poke into the indicated hole after it was illuminated (response latency) and the time from the exit from the correct hole until the entry into the reward receptacle (reward latency) were measured, whereby the latter is usually used as a compound indicator of motivation and locomotor drive[3]. In all stages, sessions lasted 30 min and were performed once daily at the same time of day and in the same box for each animal.

After surgery (see below), animals were trained until they had reached the final baseline stage (BL; 2 s SD, 5 s ITI) obtaining an accuracy >80% and an omission rate <50% in two consecutive sessions. For the last 5 d before the first imaging session, mice were accustomed to be gently fixed in the experimenter’s hand for the fixation of the miniscope before the session and were trained with dummy ‘miniscopes’ that were equal in height and weight, attached to the baseplate. Training with dummy miniscopes lasted until an accuracy >80% and an omission rate <50% in two consecutive sessions was reached again. On subsequent days, mice were trained for 3 d with an actual, tethered miniscope in distinct operant chambers set up for simultaneous imaging, followed by the first imaging session conducted also in the baseline stage. Subsequently, the imaging sessions (30 min) were repeated with different challenge conditions in the same order for every mouse (see Figure 1E). Some imaging sessions needed to be repeated due to technical failures. In between imaging sessions, mice were trained in the same testing chambers in the baseline stage with the miniscope attached. After imaging sessions with the challenge protocols were completed, some mice underwent a separate set of pharmacological experiments in the 5-CSRTT (data not shown in this manuscript), after which training in the baseline protocol and then further imaging sessions in the combined 0.8s-SD/7s-ITI challenge protocol with concomitant manipulation of value or occurrence of reward followed: Firstly, imaging was conducted under normal conditions of food-restriction and reward-delivery (baseline), secondly imaging was conducted after devaluation of reward by pre-feeding (providing 6 g of food overnight and 2 ml of milk reward 1 h before session start), thirdly, two sessions followed under conditions of normal food-restriction but omission of reward (extinction). Training in the baseline protocol without imaging was conducted before and after the devaluation session, but not between the extinction sessions.

### Surgical procedures

After the mice reached at least stage 4 (4 s SD, 5 s ITI) of the 5-CSRTT, animals were anaesthetized using isoflurane (AbbVie, G), received subcutaneous injections of analgesics (0.08 mg/kg buprenorphine, Bayer, G; 1 mg/kg meloxicam, Boehringer Ingelheim, G), and local scalp anaesthesia (200 μl of 0.025 % bupivacaine, AstraZeneca, UK) before placement in a stereotaxic frame (Kopf, US; manual digital frame, World Precision Instruments, US) with non-rupture mouse ear bars. The body temperature was stabilized using a feedback-controlled heating blanket (Harvard Apparatus, US) and the anaesthesia was maintained with 1.5 % isoflurane. The following stereotaxic coordinates (from bregma) and volumes were used for bilateral transfection of the stated areas; ACC: injection at AP +1.0, ML 0.4, DV 1.3 (300 nl) and 1.7 (200 nl); ventral mPFC: injection at AP +2.2, ML 0.3, DV 2.2 (200 nl). An AAV5-CamKIIα-GCaMP6m vector suspension (6.2*10^12^ vg/ml; University of Zürich viral vector facility; UZH-VVF, CH) was diluted down 1:1 to a final titre of 3.1*10^12^ vg/ml in 5 % sorbitol/PBS (Sigma, G) and infused using a 10 μl precision syringe (WPI, US) at an infusion rate of 100 nl/min. To minimize backflow of the virus, the needle was kept in place for 5 min at each site after infusion, and additionally for another 5 min 0.1 mm above the last infusion site. Subsequently, the wound was sutured, the mouse was allowed to recover in a temperature-controlled chamber at 36°C, and provided with mesh-food, gel-food and daily post-operative monitoring for 7 d, including application of meloxicam (Metacam, 1 mg/kg, Boehringer Ingelheim, G) on the first 3 d. The mice were kept on *ad libitum* food.

Approximately, one week after the injection of the viral construct, a gradient refractory index (GRIN) lens (Inscopix, CA, USA) was implanted. The surgery initially followed the steps described above for the virus injection and was followed by two craniotomies into the occipital and parietal bone where a screw (1 mm diameter, Precision Technologies, GB) was placed into each hole for later implant stability. A craniotomy for the lens (1 or 0.5 mm in diameter for ACC or mPFC, respectively) was made above the original infusion site. Before lowering the lens into the brain tissue, the skull was dried for better glue attachment and the lens was cleaned with 70% ethanol or 100% isopropanol. From the brain surface, the lens was held vertically by a pipette tip with negative internal pressure created by a vacuum pump and lowered by 20 µm every 30 s into the brain tissue at the original infusion site until reaching a depth of 1 mm (ACC). For mPFC, a custom-made GRIN-lens injector (“GRINjector”) was used placing the lens at 2 mm from the brain surface. Super-glue (Loctite 401, Henkel, DE) was applied to attach the lens to the skull, followed by light-curable dental adhesive (Breeze^TM^, Pentron, US). Dental cement was applied on the exposed skull with approx. 1 mm of the lens protruding out from the skull. Kwik-Sil™ (WPI, US) was applied on the lens to protect it from later mechanical damage. Post-operative care was applied as after the first surgery.

Between 2-6 wks after lens implantation, the GCaMP6m expression was checked. For this, the mouse was anaesthetized and fixed to the stereotaxic frame as described above. After removing the Kwik-Sil™ cone, the lens surface was wiped with lens tissue soaked in 70% ethanol or 100% isopropanol. The baseplate was attached to the miniscope and to the stereotactic frame using a clamp. Depending on the quality and quantity of GCaMP6m-expressing cells, the baseplate was persistently fixated to the implant by applying light-curable dental cement (Flow-It^TM^, Pentron, US) around the lens, layer by layer leaving a small (approx. 1 mm) gap below the baseplate, which was filled with 2-component adhesive (Loctite 3090, Henkel DE) for ultimate fixation. After drying, the miniscope was de-attached and a custom-made protective cap was put on the baseplate and fixated by the baseplate screw.

### Calcium imaging

Calcium imaging was done using UCLA miniscopes v3 or v4 [29], including their data acquisition (DAQ) box (https://open-ephys.org) and acquisition software (www.miniscope.org). The temporally aligned recording of behavioural events and imaging frames was achieved through pyControl [19,28], connecting the miniscope DAQ-box (https://open-ephys.org) via an input trigger GPIO SMA connector to the pyControl microcontroller board through which the start and the end of image acquisition was controlled by TTL-pulses sent to the miniscope DAQ-box. Prior to the session start, mice were equipped with the miniscope and the optimal focus was set by adjusting the focus slider of the miniscope manually (v3) or the focus electronically (v4). A thin and flexible coaxial cable (CW2040-3650SR, Cooner Wire, US) connected the miniscope to the DAQ-box for power supply, LED control, and CMOS data transmission. For some recordings a custom-made motorized commutator [30] was used to eliminate the need to manually un-twist the cable. Images were recorded at 20 fps, maximum gain, and with an excitation intensity that was adjusted for each mouse individually.

### Histology

Animals were given an over-dose of ketamine/medetomidine (≥200 mg/kg ketamine, Zoetis, G; ≥2mg/kg medetomidine, Pfizer, US) and perfused with 0.01 M phosphate-buffered saline (PBS) followed by 4 % PFA/PBS. The baseplate and the implant were carefully detached from the scull and the brain was removed and then stored in 4 % PFA/PBS overnight before placement in 20 % sucrose for dehydration before sections were cut at 100 μm thickness on a vibratome (VT1000, Leica, DE). Every second section was stained with DAPI (10^-4^ % w/v) for 30 min, washed with PBS twice and mounted on glass slides. A Leica DM6B epifluorescence microscope (Leica, DE) was used to scan the slides with a 5x objective and determine virus expression offline.

### Data analysis

#### Pre-processing of calcium traces

Single-photon imaging data for each session were pre-processed using MATLAB as described previously[17]. Each image frame was spatially down-sampled to a 400x400 pixel frame and divided by its low-pass filtered version to remove wide-field fluctuations and brightness gradients over the field of view. After band-pass filtering each frame to enhance structural features of the image to facilitate the alignment of different frames, the TurboReg algorithm [31] was used for motion correction. Each movie was temporally smoothed and temporally down-sampled from 20 Hz to 5 Hz followed by signal normalization of each image frame in units of relative changes in fluorescence, ΔF(t)/F0 = (F(t) − F0)/F0, where F0 is the mean image obtained by averaging the entire movie. For cell sorting, spatial filters corresponding to individual neurons were identified using an automated cell sorting routine based on principal and independent component analysis (PCA/ICA) [32]. Extracted spatial filters were verified as neural cells upon visual inspection based on size, morphology and the activity trace.

#### Temporal alignment of calcium traces to behavioural events

Calcium traces of each cell were z-transformed using the Matlab function *zscore* to control for variations between animals and sessions. Custom written Matlab scripts were used first to align the behavioural and the imaging data determining the timestamps for the ITI start, cue presentation, choice and outcome (referred to as epochs) of each trial in terms of frame numbers, and for labelling the trial with the corresponding choice of the mouse, i.e. correct, incorrect or premature response, or omission (referred to as response type). In each trial, the calcium signals were extracted within defined time windows from 4 s before to 7 s after the event timestamp of the onset of each epoch. The extracted traces were then averaged across trials of the same response type, thereby forming population vectors that represented each response type aligned to the onset of each epoch. Peri-event time histograms were created by plotting heat maps of the population vectors for each response type aligned to the choice onset (Figure 2). The cells were sorted based on their average peak latency. For cross-validation, heat maps were created based on the population vectors created from averaging only across even or odd trials, which were then correlated across cells within each time bin to receive a measure for the reliability of the temporal activity pattern. K-means clustering was applied using the Matlab function *kmeans* with a preset number of four clusters and the distance metric set to ‘cosine’. Thereby, the cells were grouped based on the similarity of their calcium signals, which were extracted within defined time windows relative to the onset of the respective choice (see above) and averaged across *odd* trials. Subsequently, the cells were sorted according to their cluster assignment and peak latency applying the sorting order to the calcium signals averaged across *even* trials and plotting them using peri-event time histograms (Figure 2C-D). For each cluster, the single-trial calcium signal of a corresponding exemplary cell was plotted for each response type using peri-event time histograms (Figure 2E-F).

#### Decoding analysis of population activity

Binary linear support vector machine (SVM) classifiers (Figure 3A-C and 6E-F) were trained and tested on differentiating between trials with correct responses and those with either omitted, premature or incorrect responses based on the amplitude of the calcium trace of all neurons in a FOV combined in 200 ms time-bins in a time-window from 4 s before until 7 s after the onset of each choice poke. For training the classifier, the Matlab function *fitcsvm* was used with the kernel function, box constraint, kernel scale and standardization set to ‘linear’, ‘1’, ‘auto’ and ‘false’, respectively. The dataset of every session (with a minimum of 6 events for each response type) was randomly partitioned into the training (each observation was labelled with the response type) and test set (lacking label assignment) in a 80/20 ratio, while ensuring the test set maintained balance, resulting in an imbalanced distribution within the training set for most sessions. For sessions, where the number of trials varied extensively across response types, trials from the response type with a higher number of events were randomly removed until achieving balance in the test set. For training the SVM classifier, a *balanced training set* is essential to prevent a classification bias towards the majority class (i.e. the behavioural event class with the highest number of observations in the respective data set) [35]. Using the *synthetic minority oversampling technique* (SMOTE) [36,37] on the training set, the number of observations in each event class was equalized by artificially synthesizing new samples in the minority classes (i.e. the behavioural event classes with a lower number of observations than the majority class). This algorithm randomly selects an observation from the underrepresented event class and identifies its four nearest neighbours, of which one is randomly chosen. A value is randomly picked in the Euclidean distance between the observation and the neighbour and is assigned to the new synthesized sample. The smote approach requires the number of events in the minority class to be greater than the number of set neighbours (i.e. four) and the ratio between the number of events in the majority and minority class to be less than the set number of neighbours (i.e. four). In sessions where this was not the case, events were up-sampled for the minority class using SMOTE with the number of neighbours set to the number of events in the minority class and/or trials were randomly removed from the majority class until the required conditions were met. The entire procedure of random data set partitioning, SMOTE up-sampling of the training set, and the subsequent training and testing of the decoder was repeated 100-times, thereby producing 100 averages and 100 s.e.m. values, which were averaged to yield a *grand mean* and *grand s.e.m* value representing the decoding performance based on each session. As a control for the test decoder, a second binary classifier model was established as described above, but labels of the training set in each fold within each session were shuffled prior to classifier training, thereby creating distributions that represented chance level (null distribution) [38].

Since mice mostly made an insufficient number of incorrect responses during the varITI challenges, to allow this type of decoding analysis, the data from several sessions with at least 8 incorrect responses were grouped across challenge protocols (five combined sessions, two varITI sessions, two fixed-ITI sessions) to perform a separate decoding analysis that included incorrect responses (Figure 3C).

Multiclass decoding was performed to predict into which of the five poke holes the mouse poked into during a given trial including all response types or correct responses only, using a multi-class SVM classifier [39,40] with the same approach as described above (Figure 4D). The linear SVM multi-classifier was trained using the matlab function *fitcecoc* with the kernel function, box constraint, kernel scale and standardization set to ‘linear’, ‘1’, ‘auto’ and ‘off’, respectively. Additionally, the option coding was set to ‘onevsall’; the one-vs-all strategy performs a separate binary classification for each class in the dataset (i.e. in total four) treating it as the positive class, whereas all other classes combined are treated as the negative class. Testing is performed by independently applying every sample from the test data set on each trained binary classifier yielding confidence values with the highest selecting the predicted class for this sample.

#### Encoding analysis of the modulation of neural activity by behavioural events

Linear regression models were created to predict the calcium signal in 200 ms timebins in a time window from 4 s before until 7 s after the onset of the choice epoch for each individual neuron (Figure 5 and 7). Regularized linear regression was performed using the Matlab function *lasso* applying L1 (lasso) regularization with 10-fold cross-validation to find the optimal regularization strength λ that minimizes the loss. Binary predictors were used to code for the presence of a poke (active poke vs. omission), the spatial poke identity (four one-hot predictors corresponding to left-right directionality, right poke discrimination, left poke discrimination and middle poke discrimination), and correct (rewarded) responses (vs. erroneous and omitted responses combined) in each trial (see Figure 5A for predictor matrix). To test how much of the variance of the activity of individual neurons at every time bin could be explained by each predictor, the coefficient of partial determination (CPD) was calculated, measuring how much further the predictor contributed to the explanation of the full regression model [21,33,34]. CPDs were determined for each predictor by subtracting the mean squared error of the full linear regression model from the mean squared error of the reduced regression model, where the predictor for the specific event type in question was removed. The CPD for predictor *i* is defined as:

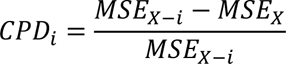

where *MSE_X-i_* is the mean squared error in a regression model that includes all of the relevant predictor variables except *i*, and *MSE_X_* is the mean squared error in a regression model that includes all of the relevant predictor variables. To compute the CPD for spatial poke identity, all of the four spatial one-hot predictors were removed together.

For the devaluation and extinction conditions (Figure 7), which led to reduced numbers of correct responses, pseudo-randomly chosen events from the baseline (control) condition were selected to roughly match the number of the respective experimental condition for each poke-hole and response type, to ensure that CPDs in the baseline session are not simply higher because of a larger number of trials or responses used for their calculation. This down-sampling was repeated 100 times and the CPDs of the respective predictors averaged across iterations before plotting.

### Statistics

Behavioural data was analysed using Matlab (R2019a, The MathWorks Inc, USA) and only two-sided tests were used. 5-CSRTT performance during calcium imaging sessions (Figure 1D-E, Figure 6A-D) was analysed using an ANOVA involving the task paradigm as between-subject independent variable and one of the behavioural parameters as dependent variable. In case of a significant effect of task paradigm, Dunnett’s post-hoc tests were conducted between the baseline and any other challenge. Decoding accuracies (Figure 3) were statistically compared using repeated-measures ANOVA with the time-bin and epoch type variable as within-subject factors. A Dunn-Sidak-test was used for post-hoc testing. Comparisons against accuracies of control classifiers (trained with shuffled labels, performing at chance level) in decoding analyses or against 0% CPD in encoding analyses have been done with paired-sample or one-sample *t*-tests, respectively, with Benjamini-Hochberg corrections for the repeated testing in each time interval. All applied statistical tests are stated in the corresponding figure legends. All bar and line graphs display mean ± s.e.m. or data from individual mice, as indicated.

## Data availability

All raw data can be obtained from the corresponding author upon reasonable request. Scripts of all task files applied in custom-made operant boxes can be obtained from https://github.com/KaetzelLab/Operant-Box-Code and design files for such operant boxes are deposited at https://github.com/KaetzelLab/Operant-Box-Design-Files.

## Code availability

Analysis scripts are available from GitHub at https://github.com/martinjendryka/Jendryka_et_al_ACC_imaging_5CSRTT.git.

## Acknowledgments

We thank Stefanie Schulz (Ulm University) for assistance with histology. <u>Funding</u>: This work was funded by the Boehringer Ingelheim-Ulm University (BIU) Center (TPN010, to B.L., A.P., D.K.), the Else-Kroener-Fresenius/German-Scholars-Organization Programme for excellent medical scientists from abroad (GSO/EKFS 12; to D.K.), the DFG (KA 4594/2-1; to D.K.), and the Alfred-Krupp Foundation (to B.L.).

## Author Contributions

M.M.J., B.L., A.P., T.A. and D.K. designed the study. M.J. and U.L. conducted behavioural experiments. M.M.J. conducted surgeries. S.K.T.K., T.A., and D.K. developed pyOS-5 operant box hardware and software; S.K.T.K. programmed operant box task protocols and integration of miniscope recordings. B.F.G. and H.D. provided assistance with manufacturing and usage of UCLA v3 miniscopes. B.F.G., B.L. and D.K. provided essential resources. M.M.J. analysed the data with advise from B.F.G., T.A. and D.K.. M.M.J. and D.K. wrote the manuscript, which was revised by all authors.

