## Supplementary Information for "Excitatory neurons of the anterior cingulate cortex encode chosen actions and their outcomes rather than cognitive state"

### Supplementary Figures

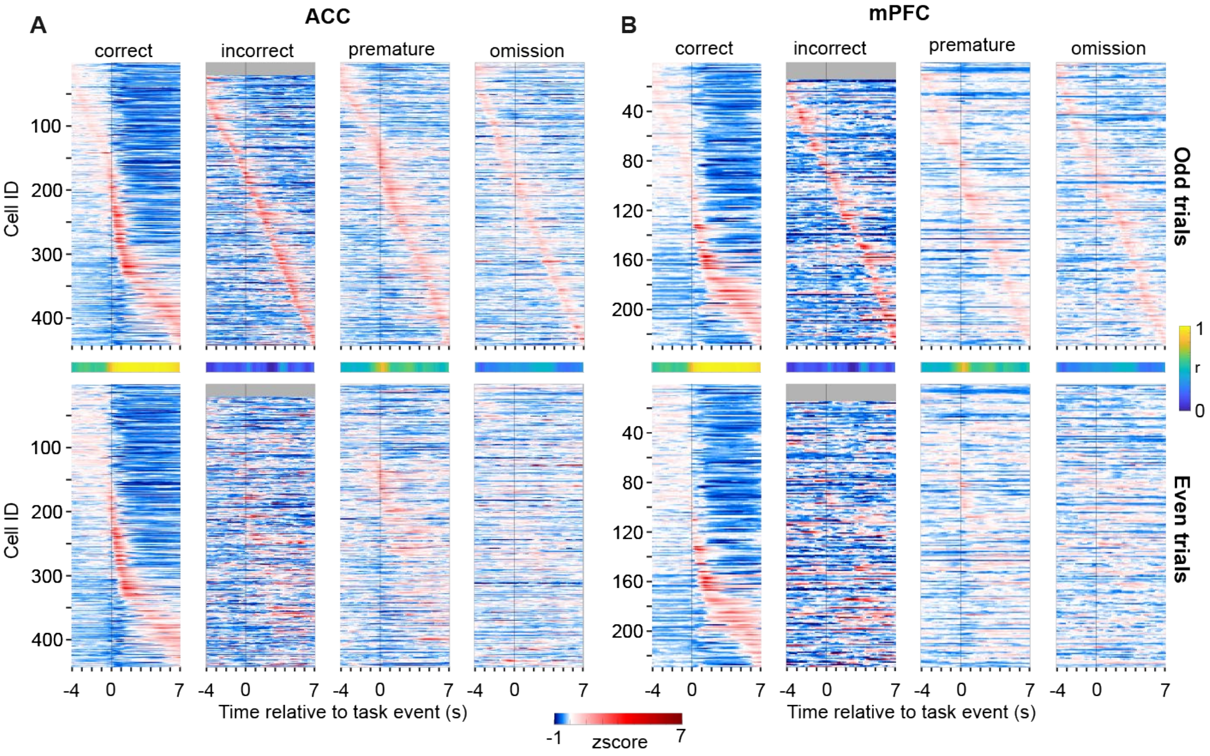


**Supplementary Figure 1. Detection of event-locked activity of individual neurons in the variable ITI challenge**. Exemplification of cross-validation of time-locked average activity using separate averages for *odd* and *even* trials. (**A**) *Top*: Average z-scored calcium activity in individual ACC neurons time-locked to the onset of the behavioural event stated above each sub-panel, shown for -4 - +7 s around the event and sorted according to the time of occurrence of the maximum amplitude. Averaging was done across the trials with *odd* order numbers. *Bottom*: Similar display as in corresponding top panels, but the average was calculated from trials with *even* order numbers whereas the top-to-bottom order determined from odd trials (as shown in top panels) was maintained. *Middle*: Pearson correlations between the top and the bottom panel at each time point. Note that temporal order of peaks is maintained for correct and premature responses with resulting high correlations, but not for incorrect choices and omissions. *N* = 12 animals and 443 cells. (**B**) Same display and analysis as in (A) but for all neurons recorded in mPFC. *N* = 6 animals and 229 cells.

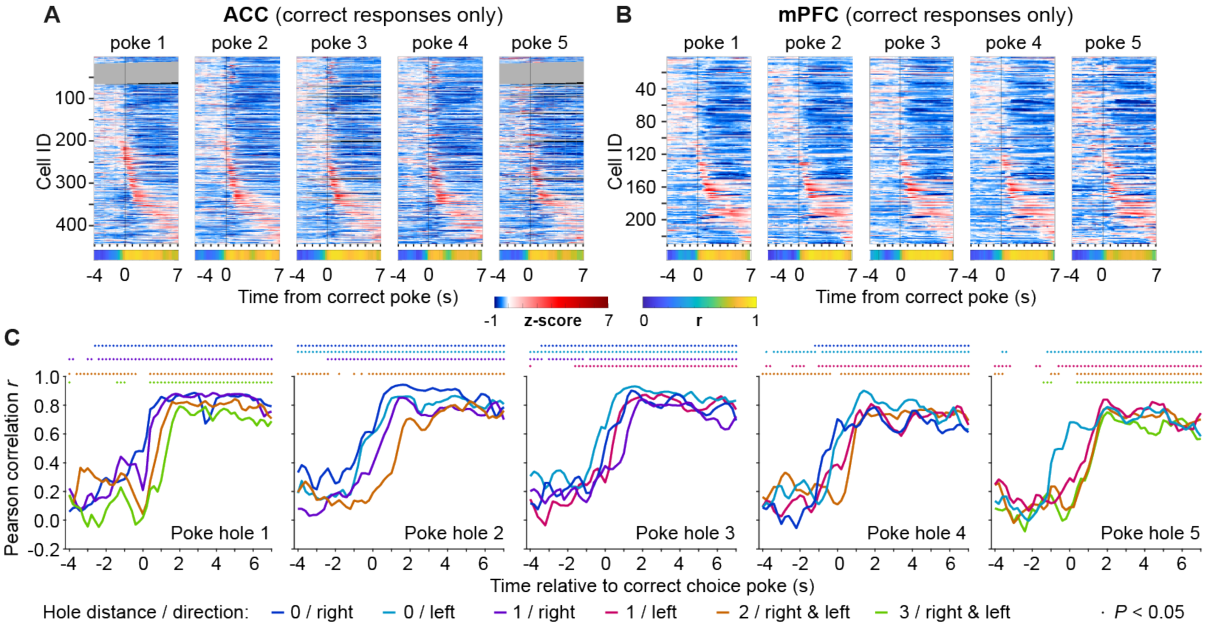


**Supplementary Figure 2. Activity in the ACC represents spatial action selection.** (**A-B**) Same data as in Figure 2A-B (correct responses) but separated by poke-hole. *Top:* Average z-scored calcium activity in individual ACC (A) or mPFC (B) neurons time-locked to the correct choice-poke, shown for -4 - +7 s around the event. For cross-validation, averaging was done across the *even* trials only and the cells were sorted according to the average peak latency across the *odd* trials of poke 1. Grey lines indicate sessions in which the given hole was not poked into. *Bottom:* Pearson correlations between the averaged z-scored activity of the *odd* and *even* trials at each time point. Note that the qualitative similarity to the pattern of poke 1 gets reduced the further away the poke-hole is. (**C**) Based on the data shown in (A), Pearson correlations between response patterns of pairs of poke holes, coded in colour according to the distance between the holes and direction relative to the reference hole named in the lower right corner of each sub-panel. Note that correlations have been calculated only across cells for which the Pearson correlation of activity patterns between odd and even correct trials was above the median correlation to ensure that cells used for this analysis actually display time-locked response-related activity. The dots at the top indicate significance ($P$ < 0.05) of the correlation coefficient plotted in the matching colour in the sub-panel below.

**A** **Orthogonal predictor matrix**

**Encoded behavioural event**

| Predictor | poke 1 | poke 2 | poke 3 | poke 4 | poke 5 | omission |
|---|---|---|---|---|---|---|
| active poke / omission | 1 | 1 | 1 | 1 | 1 | -1 |
| left-right directionality | 1 | 1 | 0 | -1 | -1 | 0 |
| poke discrimination right | 0 | 0 | 0 | 1 | -1 | 0 |
| poke discrimination left | 1 | -1 | 0 | 0 | 0 | 0 |
| poke discrimination middle | -0.25 | -0.25 | 1 | -0.25 | -0.25 | 0 |

| | rewarded trial | non-rewarded trial |
|---|---|---|
| rewarded (correct) response | 1 | -1 |

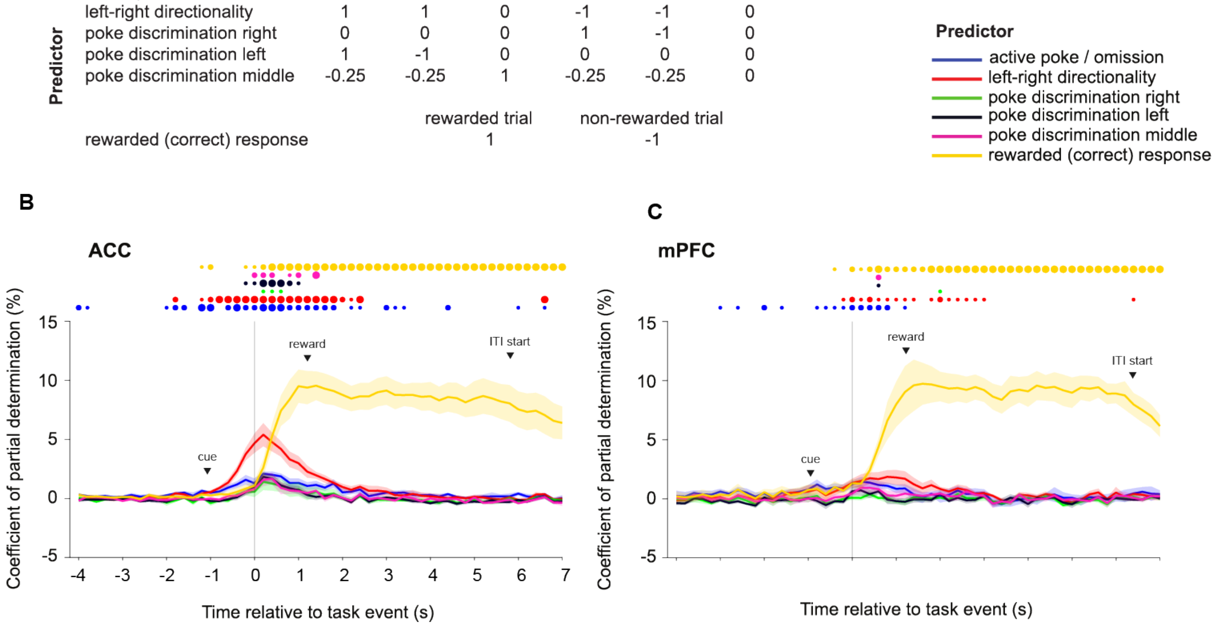


**Supplementary Figure 3. Encoding of poking action and reward in the population activity of the ACC and mPFC.** Analysis as in main Figure 5, but with individual removal of spatial predictors. (**A**) Orthogonal predictor matrix designed to indicate the representation of the poke (1: poke in either hole, 0: omission), the poke directionality (1: poke in left holes 1 or 2; -1: poke in right holes 4 or 5; 0: poke in middle hole 3 or omission), right poke discrimination (1: poke in hole 4; -1: poke in hole 5; 0: poke in holes 1,2 and 3 and omission), left poke discrimination (1: poke in hole 1; -1: poke in hole 2; 0: poke in holes 3,4 and 5 and omission), middle poke discrimination (1: poke in hole 3; -0.25: poke in holes 1,2,4,5; 0: omission) and reward (1: rewarded; 0: not rewarded). (**B-C**) Coefficient of partial determination (CPD) averaged across cells recorded in ACC (B, N = 11 mice) and mPFC (C; N = 6 mice). Time bins where CPDs for a given event were significantly higher than zero after cross-validated linear regression are indicated with a dot at the top of each panel, colour-coded for the respective predictor (one sample *t*-test with Benjamini-Hochberg post-hoc correction). CPDs were determined for each event type by subtracting the sum squared errors of the full linear regression model (incorporating every event type as predictor) from the sum squared error of the reduced regression model where one predictor (corresponding to the event type for which the population activity should be explained) was removed.

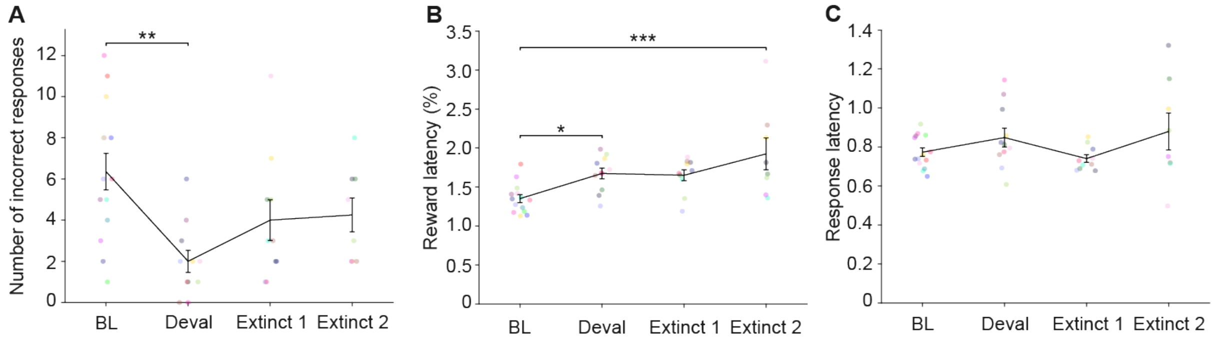


**Supplementary Figure 4. Behavioural performance during devaluation and extinction.** (**A-C**) Measures of task engagement and performance in the 5-CSRTT (as indicated on y-axes) during training sessions in the 0.8s-SD/7s-ITI combined challenge at normal reward conditions (baseline, BL), after devaluation of reward (Deval), or with omission of reward (extinction; Extinct 1/2), as indicated on x-axes with tethered miniscope. Dots indicate individual animals coded by colour, bars show mean ± s.e.m.. Asterisks represent Dunnett pairwise post-hoc test comparing BL condition against the other conditions after RM-ANOVA. * $P < 0.05$; ** $P < 0.01$; *** $P < 0.001$.

#### Supplementary Tables

| | | ACC | | | mPFC | | |
|---|---|---|---|---|---|---|---|
| **challenge** | **Figure** | **classlabel 1** | **classlabel 2** | **N** | **classlabel 1** | **classlabel 2** | **N** |
| **varITI** | - | correct | incorrect | 2 | correct | incorrect | 1 |
| | **3A-B** | **correct** | **omission** | **11** | **correct** | **omission** | **6** |
| | **3A-B** | **correct** | **premature** | **11** | **correct** | **premature** | **6** |
| **mixed** | **3C** | **correct** | **incorrect** | **6** | correct | incorrect | 3 |
| | - | correct | omission | 6 | correct | omission | 3 |
| | - | correct | premature | 6 | correct | premature | 3 |
| **combined: baseline** | - | correct | incorrect | 5 | correct | incorrect | 3 |
| | **7A** | **correct** | **omission** | **10** | correct | omission | 4 |
| | **7B** | **correct** | **premature** | **9** | correct | premature | 4 |
| **combined: devaluation** | - | correct | incorrect | 1 | correct | incorrect | 0 |
| | **7A** | **correct** | **omission** | **5** | correct | omission | 5 |
| | **7B** | **correct** | **premature** | **3** | correct | premature | 4 |
| **combined: extinction1** | - | correct | incorrect | 1 | correct | incorrect | 1 |
| | **7A** | **correct** | **omission** | **5** | correct | omission | 4 |
| | **7B** | **correct** | **premature** | **5** | correct | premature | 4 |
| **combined: extinction2** | - | correct | incorrect | 2 | correct | incorrect | 1 |
| | **7A** | **correct** | **omission** | **5** | correct | omission | 2 |
| | **7B** | **correct** | **premature** | **4** | correct | premature | 2 |

**Supplementary Table 1. Number of animals contributing to classifiers.** *N*-numbers of animals contributing data to the classifier results shown in Figures 4 and 8 are stated in dependence on the challenge (experiment) and the response type that the classifier discriminated. Numbers for data that is displayed are in bold, contrast that were not calculated because of low N-number are shown for information in non-bold font. The reduction of N-numbers compared to the number of mice that participated in experiments (ACC: 12, mPFC: 6) results primarily from low (< 6) numbers obtained for one response type, but in rare cases also from a strong imbalance of response numbers, that cannot be compensated for by the smote-approach, or from technical failures in experiment execution or in data acquisition.
